# Stress-induced dysfunction of neurovascular astrocytes contributes to sex-specific behavioral deficits

**DOI:** 10.1101/2024.05.14.594147

**Authors:** Justin L. Bollinger, Shobha Johnsamuel, Lauren L. Vollmer, Alexander M. Kuhn, Eric S. Wohleb

## Abstract

Astrocytes form an integral component of the neurovascular unit, ensheathing brain blood vessels with projections high in aquaporin-4 (AQP4) expression. These AQP4-rich projections facilitate interaction between the vascular endothelium, astrocytes, and neurons, and help stabilize vascular morphology. Studies using preclinical models of psychological stress and post-mortem tissue from patients with major depressive disorder (MDD) have reported reductions in AQP4, loss of astrocytic structures, and vascular impairment in the prefrontal cortex (PFC). Though compelling, the role of AQP4 in mediating stress-induced alterations in blood vessel function and behavior remains unclear. Here, we address this, alongside potential sex differences in chronic unpredictable stress (CUS) effects on astrocyte phenotype, blood-brain barrier integrity, and behavior. CUS led to pronounced shifts in stress-coping behavior and working memory deficits in male –but not female– mice. Following behavioral testing, astrocytes from the frontal cortex were isolated for gene expression analyses. We found that CUS increased various transcripts associated with blood vessel maintenance in astrocytes from males, but either had no effect on-or decreased-these genes in females. Furthermore, CUS caused a reduction in vascular-localized AQP4 and elevated extravasation of a small molecule fluorescent reporter (Dextran) in the PFC in males but not females. Studies showed that knockdown of AQP4 in the PFC in males is sufficient to disrupt astrocyte phenotype and increase behavioral susceptibility to a sub-chronic stressor. Collectively, these findings provide initial evidence that sex-specific alterations in astrocyte phenotype and neurovascular integrity in the PFC contribute to behavioral and cognitive consequences following chronic stress.

## Introduction

Astrocytes form an integral component of the neurovascular unit, ensheathing brain blood vessels with endfoot projections rich in aquaporin-4 (AQP4) [1–5]. These AQP4-endfeet play a critical role in water and ion exchange, and facilitate interaction between astrocytes, the vascular endothelium, and various synaptic components [6]. For instance, genetic ablation of AQP4 in mice reduces astrocyte glutamate uptake, impairs glucose trafficking, destabilizes the vasculature, and disrupts glymphatic flux [7–12]. Constitutive, brain-wide knockout of AQP4 has even been shown to shift aspects of behavior, including spatial- and associative fear memory [7, 10].

Studies using preclinical models of chronic psychological stress and post-mortem tissue from patients with major depressive disorder (MDD) have reported reductions in AQP4, loss of perivascular astrocytic processes, and atrophy of astrocytes in the prefrontal cortex (PFC) [9, 13–19]. Some of these structural changes in astrocytes (i.e., reductions in GFAP+ material) are associated with loss of synapses in the PFC, anhedonia-like behavior, and impaired active avoidance learning in preclinical models [20–22]. Alongside pronounced astrocyte dystrophy, recent studies indicate disruption of the neurovasculature in stress. This includes loss of tight junction proteins, reduced cerebral blood flow, and heightened blood-brain barrier (BBB) permeability in select brain regions [9, 23–28]. Together, these findings suggest that stress-induced dysregulation of astrocytes at the PFC neurovascular interface leads to behavioral and cognitive consequences.

To test this directly, we carried out experiments examining the role of astrocyte AQP4 in regulating stress effects on PFC architecture and function. We first characterized chronic unpredictable stress (CUS) effects on astrocyte phenotype, astrocyte-blood vessel interaction, BBB integrity, and behavior in both male and female mice. Our data indicate basal sex differences in astrocyte structure and blood vessel coverage in the PFC, alongside opposite patterns of stress-induced astrocyte remodeling in males and females. Stress effects on astrocytes were also associated with divergent patterns of cognitive and behavioral dysfunction in males and females. In subsequent experiments, we knocked down *Aqp4* in the PFC using an shRNA-mediated approach. Loss of prefrontal AQP4 altered the molecular- and structural phenotype of astrocytes in this region and induced behavioral susceptibility to stress in male mice. Collectively, our findings support a sex-specific role for neurovascular astrocytes in regulating chronic stress effects on behavior.

## Materials and Methods

### Chronic unpredictable stress (CUS, 14 days) and sub-CUS (7 days)

Chronic unpredictable stress was performed as previously described [29]. In brief, mice were exposed to two random intermittent stressors per day for either 7 (sub-CUS) or 14 days (CUS). Stressors included: cage rotation, isolation, restraint, radio noise, food or water deprivation, light on overnight, light off during day, rat odor, stroboscope overnight, crowding, wet bedding, no bedding, or tilted cage. This stress protocol produces reliable increases in circulating corticosterone, adrenal hypertrophy, and reduced weight gain. A 7-day, sub-CUS protocol was selected based on previous studies showing limited stress-induced deficits in behavior, thus allowing for the assessment of stress susceptibility [30–32]. To validate this manipulation, body weight was measured at the start and end of CUS and sub-CUS, with percent weight gain calculated (Fig S1).

### Short-hairpin RNA (shRNA) and viral preparation

An shRNA targeting the following *Aqp4* transcript sequence (NM_009700: 5’-GCACACGAAAGATCAGCATCG-3’) was designed using VectorBuilder software [12]. This *Aqp4* shRNA (5’-GCACACGAAAGATCAGCATCG-CTCGAG-CGATGCTGATCTTTCGTGTGC-3’) or a scrambled control shRNA was integrated into a pTagBFP2-shRNA construct driven by the cytomegalovirus (CMV) promoter with shRNA sequences driven by the U6 promoter. Databases indicate that *Aqp4* is selectively expressed by astrocytes in the cortex, so these promoters were chosen to maximize construct expression and transcript knockdown [33]. Constructs were then packaged into adeno-associated virus 5 (AAV5) following VectorBuilder’s standard procedures.

### Stereotactic surgery and viral infusions

Mice were deeply anesthetized with ketamine:xylazine (90 mg/kg:20 mg/kg) followed by stereotactic surgery. Virus containing either a scrambled shRNA or *Aqp4*-knockdown shRNA (2L10^13^ GC/ml) was bilaterally infused (0.8 µl; 0.1 µl/min) into the mPFC (AP: +2.0mm, ML: ±0.2mm, DV: −2.8mm). Syringes were left in place for 5 min after infusion to limit viral spreading in the needle tract. Mice received daily injection of meloxicam (5 mg/kg, i.p.) over 3 days for pain relief. Mice were handled intermittently over three weeks to allow for surgical recovery and viral infection/expression.

### Behavior and cognitive tests

Open field test (OFT), forced swim test (FST), and temporal object recognition testing (TOR) were conducted as previously described [22, 34]. A cognitive T-maze task was used to assess spatial reference memory and short-term memory, as adapted from previous T- and Y-maze testing paradigms [35–38]. The mPFC contributes to stress coping behavior in the FST and working memory function as assessed in the TOR and cognitive T-maze task, making these behavioral assays particularly relevant to the neurobiological outcomes examined in this study [39, 40]. The open field test was used to assess animal mobility in a novel context and thigmotaxis. All behavioral tests were performed in a normally lit room (white light, 300 lux), between 0800 and 1200 hours. Mice were allowed to habituate to this room for 30 min prior to testing. For the OFT, individual mice were placed in an empty, white Plexiglas™ arena (40 × 40 × 30 cm). The center of the OFT was defined as the innermost zone (8 × 8 cm). Activity was recorded for 10 min and both mobility and the total time spent in the center zone were assessed using Noldus EthoVision XT 13. For the FST, mice were placed in a 2-liter beaker of water (24°+/-1°C) for 8 min and time spent immobile (2-6 min phase) was measured. Only animals undergoing CUS were exposed to the FST, with behavior assessed on day 0 (prior to CUS) and day 11. In the first trial phase of the TOR, mice were placed in a plastic arena with two plastic Lego™ trees secured to the bottom of the arena. For each phase of the TOR, mice were given 5 min to explore the arena and objects. After a 30 min latency, mice were placed in the same arena with Lego™ blocks (second trial phase). An hour after the second trial phase, mice were placed in the arena with one block and one tree, counterbalanced to prevent bias (test phase). The time spent exploring each of these objects was measured and a discrimination index was calculated (difference between time spent exploring the tree or the block divided by the total time spent exploring the tree and the block). In the trial phase of the T-maze task, mice were placed in the distal end of the start arm with one choice arm closed. To prevent choice bias, the closed arm (left or right) was counterbalanced across subjects. Extra-maze visual cues were fixed above each choice arm. Animals were given 8 min to explore the T-maze in this configuration, after which they were placed back in their home cage. Following a 40 min interval, mice were returned to the start arm of the T-maze with both choice arms now open (test phase). Mice were allowed to freely explore the maze for 5 min. EthoVision was used to calculate the total time spent exploring each choice arm (i.e., familiar arm vs. novel arm). In addition to these metrics, the number and sequence of arm entries was recorded. Spontaneous alternation (%) was calculated as the number of consecutive arm entries divided by the total number of alternations minus 2. For CUS studies, behavioral testing occurred on day 0 (pre-stress FST), 11 (post-stress FST), 12 (OFT), 13 (TOR), and day 14 (T-maze). For sub-CUS studies, testing occurred on day 5 (OFT), 6 (TOR), and 7 (T-maze). All testing apparatuses were cleaned with 70% EtOH between trials. All observer scored metrics were obtained by a trained researcher blinded to animal condition.

### Percoll gradient enrichment of astrocytes and fluorescence activated cell sorting (FACS)

For these studies, mice were euthanized by rapid cervical dislocation followed immediately by brain extraction. Whole brains were then split by making a transverse cut (ventral-to-dorsal) at the optic chiasm. Cortex rostral to this split was enriched for the PFC by dissecting out all structures below the forceps minor/corpus callosum followed by removal of ventral cortex (structures below agranular insular cortex). Dissected frontal cortex (cortex rostral to the optic chiasm) was minced and immersed in an enzyme incubation solution (1 mL) containing 200 U papain (Worthington Biochemical, #LS003126), 0.5 mM EDTA (Invitrogen, #AM9260G), and 5 μg actinomycin-D (Millipore Sigma, #A1410) in 1X HBSS/PBS (1:1, pH 7.4±0.2, ThermoFisher, #14065056) [41]. Samples were incubated at 37°C for 10 min (water bath), followed by gentle trituration (p-1000 pipette), an additional 5 min incubation step, and a final round of gentle trituration (p-200 pipette). Outside of this incubation period, samples were stored on ice to reduce isolation associated shifts in cell phenotype. Homogenates were then suspended in an enzyme inhibitor solution containing ovomucoid (15 mg/mL; Worthington Biochemical, #LS003086) and bovine serum albumin (BSA, 15 mg/mL; Sigma-Aldrich, #A4161), followed by centrifugation at 800g for 8 min. Supernatant was removed and cell pellets were re-suspended in 30% isotonic Percoll (GE Healthcare, #17089102). Cell suspensions were centrifuged for 20 min at 2000g. Supernatant was removed, and cell pellets were washed and re-suspended. Fc receptors were blocked with an anti-CD16/CD32 antibody (BioLegend, #553141) for 10 mins. Cells were then incubated with conjugated antibodies (FITC-CD11b, BD Biosciences, #557396; PE-Recombinant-ACSA2, Miltenyi Biotec, #130-116-244) for 45 min at 4°C. Cells were washed and re-suspended in fluorescence activated cell sorting (FACS) buffer. An average of 26,333±1445 CD11b-/ACSA2+ astrocytes were isolated per sample using a BioRad S3e four-color cytometer/cell sorter. Data were analyzed using FlowJo software.

### RNA extraction, cDNA preparation, and quantitative real-time polymerase chain reaction (qPCR)

RNA was extracted from whole brain regions (PFC and somatosensory cortex; SSCTX) using TRIzol Reagent according to manufacturer’s protocol (Invitrogen). RNA was extracted from astrocytes using a Single Cell RNA purification kit in combination with DNAse treatment (Norgen Biotek Corp., #51800). Samples were reverse transcribed using a High-Capacity cDNA Reverse Transcription Kit (ThermoFisher Scientific, #4368814). Quantitative real-time PCR was conducted using a QuantStudio 5 Real-Time PCR System (ThermoFisher Scientific). All samples were run in triplicate. Cycle threshold (CT) was determined for all genes and normalized to the geometric mean of *Gapdh* and *Tbp*. Fold change was standardized to control males using the 2^-ΔΔCT^ method. Primer sequences are listed in Table S1.

### Dextran extravasation, blood vessel labeling, and immunohistology

Approximately 4 h after the final stressor, mice were briefly anesthetized with isoflurane followed by an intravenous injection of Alexa Fluor 488-conjugated dextran (10,000 MW, 100 µl; Invitrogen, #D22910). This was allowed to circulate for 30 min during recovery. Mice were then administered a lethal dose of sedative (sodium pentobarbital) followed by an intravenous injection of DyLight 594-conjugated tomato lectin (100 µl; Vector Laboratories, #DL-1177-1). Mice that received stereotactic surgery were only given injections of tomato lectin (no prior anesthesia/administration of dextran). After 5 min, mice were transcardially perfused with PBS followed by 4% paraformaldehyde (PFA). Brains were post-fixed in 4% PFA for 24 h and incubated in 30% sucrose until cryoprotected. Brains were then rapidly frozen and sectioned on a Leica CM2050 S cryostat (40 µm). Free-floating sections containing the PFC were selected for analysis. Sections used for analysis of dextran extravasation were washed for 5 min in PBS, immediately mounted on gel-coated slides, and coverslipped with Fluoromount-G (ThermoFisher Scientific, #00-4958-02). For immunohistology, sections were washed, blocked in 1% bovine serum albumin (BSA; Fisher Scientific, #BP9703) with 5% normal donkey serum (NDS; EMD Milipore, #S30-100ML) for 1 h at room temperature, washed, and then incubated with primary antibody: rabbit anti-AQP4 (1:500, Millipore-Sigma, #AB2218), rat anti-GFAP (1:1000, Invitrogen, #13-0300), or rabbit anti-ALDH1L1 (1:500, Cell Signaling Technologies, #85828), overnight at 4°C. Sections were then washed and incubated with conjugated secondary antibody overnight at 4°C (1:1000, Invitrogen; Alexa Fluor 488 or Alexa Fluor 647). Sections were subsequently washed, mounted, and coverslipped.

### Quantitative immunofluorescence

Confocal images were obtained on a Nikon C2+ microscope interfaced with a Nikon C2si+ camera. Confocal images were captured from adjacent brain sections in both hemispheres of the mPFC or the S1J subregion of the SSCTX (3-4 sections/sample spanning the rostral-caudal axis, 2.10 – 1.34 mm Bregma). For analysis of GFAP+ material, ALDH1L1+ cell counts, neurovascular AQP4+ material, tomato lectin+ vascular morphology, and dextran extravasation, tissue sections were imaged using a 20× objective (NA: 0.95, z-stack: 0.6 μm, image size: 1024×1024). Individual ALDH1L1+ cell counts were obtained by hand and cell density was calculated (cells per mm^2^). As a proxy for astrocyte structure, GFAP+ images were thresholded and total area was recorded (μm^2^). The amount of GFAP covering tomato lectin+ vessels was determined by first subtracting the vascular area from GFAP+ images using ImageJ’s Image Calculator function (NIH). This GFAP+ material was thresholded and these values were subtracted from the total GFAP+ area as previously calculated (μm^2^). Neurovascular AQP4+ material and tomato lectin+ vascular structures were thresholded and total area for each was recorded (μm^2^). To examine relative coverage of the neurovasculature by astrocytic endfeet, AQP4+ area was divided by vascular area. For analysis of dextran extravasation, sites were identified and counted for each image. These sites were defined as a concentrated 488-signal occurring directly adjacent to a blood vessel. Vessels (546-channel) were then subtracted from dextran images (488-channel) using ImageJ’s Image Calculator function (NIH). An 80 × 80-pixel region of interest was placed over each dextran extravasation site and fluorescence intensity was measured within these bounds (integrated density, A.U.), this was then standardized to control males.

### Statistical analysis

Data were analyzed using GraphPad Prism 9 (La Jolla, California). Global main effects and interactions were determined using two-way ANOVA (between-subjects; stress×sex, stress×virus). A two-way repeated measures ANOVA was used for analysis of the FST (stress×sex). Analysis of arm preference in the T-maze was done separately for males and females and animals treated with different viral constructs using two-way ANOVA (arm×stress). Following a significant ANOVA finding (main effect or interaction, *p* ≤.05), a series of hypothesis-informed, planned comparisons were conducted using Sidak’s multiple comparisons test. For studies using males and females, the following comparisons were made: male control vs female control, male control vs male CUS, male control vs female CUS, female control vs female CUS. For *Aqp4* knockdown studies, the following comparisons were made: scrambled shRNA control vs *Aqp4* shRNA control, scrambled shRNA control vs scrambled shRNA sub-CUS, scrambled shRNA control vs *Aqp4* shRNA CUS, *Aqp4* shRNA control vs *Aqp4* shRNA CUS. Pearson correlation coefficients were computed and analyzed for select variables. The number of animals examined in each analysis is noted in Table S2, with complete omnibus statistics and pairwise comparisons (including confidence intervals) reported in Table S3 and Table S4, respectively.

## Results

### Exposure to chronic stress induces sex-specific alterations in cognition and behavior

To examine the effects of CUS on cognitive and behavioral function in both male and female mice, CUS-animals were exposed to the FST prior to (d0)- and during stress (d11). All mice were then assessed in the OFT (d12), TOR (d13), and T-maze task (d14). CUS increased immobility in the FST (F_(1,38)_=12.70, *p*=0.001), with planned comparisons indicating heightened immobility in male mice only (*p*=0.024; Fig 1B). Exposure to CUS had no effect on the total distance moved in the OFT (Fig 1C) but decreased the amount of time spent exploring the center zone of the open field (F_(1,66)_=3.776, *p*=0.056) in males (*p*=0.037; Fig 1D). CUS had no effect on the total amount of time spent exploring objects in the TOR (Fig 1E), but reduced object discrimination in this task (F_(1,75)_=23.77, *p*<0.0001; Fig 1F). Planned comparisons indicate that this was significant in male, but not female, mice (*p*<0.0001). During the test phase of the T-maze task (Fig 1G), both male (F_(1,76)_=71.58, *p*<0.0001) and female mice (F_(1,74)_=30.30, *p*<0.0001) spent more time in the previously unexplored (novel) arm, regardless of stress (Fig 1H). This would suggest intact spatial recognition or novelty seeking in these mice [38]. Nonetheless, CUS increased the total number of alternations between arms (F_(1,75)_=4.988, *p*=0.028; Fig 1I) and impaired spontaneous alternation behavior (F_(1,75)_=7.717, *p*=0.007; Fig 1J) in males only (*p*=0.025 and *p*=0.039, respectively). Together, these data indicate sex-specific stress effects on cognition and behavior, with prominent CUS-induced shifts in PFC-mediated working memory tasks in male mice.

**Figure 1.**
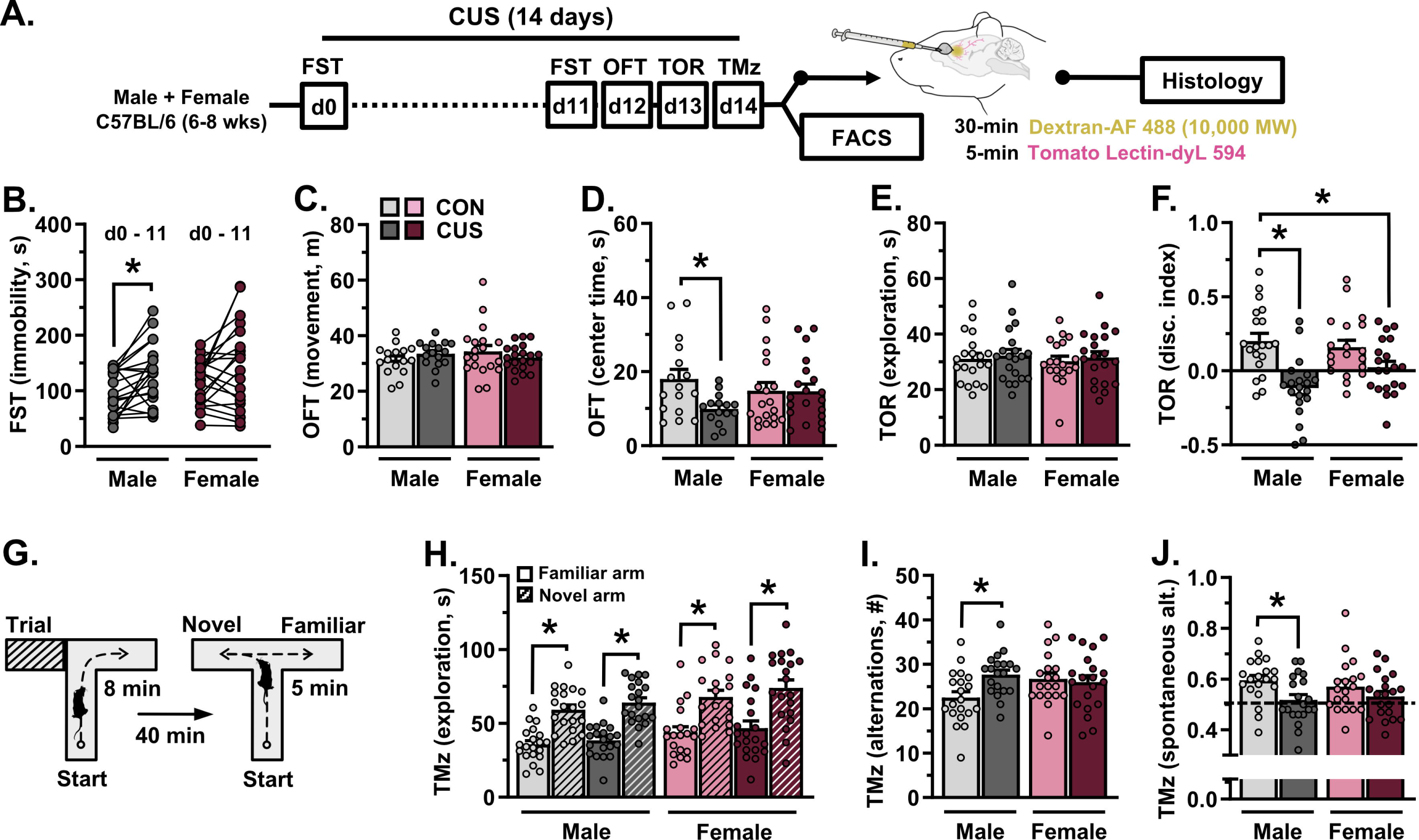
Chronic unpredictable stress (14-days) induces sex-specific patterns of cognitive and behavioral dysfunction. **A)** Adult male and female mice were exposed to 14-days of CUS or were left unstressed. All animals were subjected to behavioral testing (n = 15-20/group). In one cohort, mice received intravenous injections of Alexa Fluor 488-conjugated dextran and tomato lectin, after which brains were processed for confocal microscopy (n = 7-8/group). In a separate cohort, astrocytes were isolated via FACS and astrocyte gene expression was analyzed (n = 8-12/group). **B)** Average time spent immobile in the forced swim test (FST) prior to- and on CUS day 11. **C)** Total amount of distance traveled in the open field test (OFT). **D)** Time spent in the center during the OFT. **E)** Total amount of time spent exploring objects in the test phase of the temporal object recognition task (TOR). **F)** Discrimination index in the TOR. **G)** Schematic of the cognitive T-maze task. **H)** Time spent exploring both the novel and familiar arm in the T-maze task. **I)** Total number of alternations in the T-maze task. **J)** Proportion of spontaneous alternations in the T-maze task. Bars represent mean ± S.E.M. * p<0.05 planned comparison indicated (Sidak’s test).

### Chronic stress impacts the molecular phenotype of PFC astrocytes in males and females, broadly engages astrocyte-blood vessel communication pathways in males

To characterize sex differences in- and stress effects on the molecular phenotype of astrocytes, a PFC-enriched sample of cortex was dissected out from a subset of behaviorally tested mice (d15), astrocytes were isolated from this sample using FACS, mRNA was extracted, and astrocyte gene expression was examined (Fig 2A). CUS decreased expression of the astrocyte intermediate filament genes *Gfap* and *Vim* in both male and female mice (F_(1,35)_=13.45, *p*<0.001; F_(1,35)_=4.805, *p*=0.035), and diminished levels of *S100b* (Ca^2+^ binding factor; F_(1,35)_=8.462, *p*=0.006) mRNA in males only (*p*=0.013; Fig 2B). Expression of *Mertk*, a phagocytic receptor found on astrocytes, was increased following CUS in males (F_(1,35)_=4.866, *p*=0.034). Exposure to CUS increased expression of the angiogenesis factor *Fgf2* in both males and females (F_(1,33)_=19.83, *p*<0.0001), and selectively elevated levels of *Agt* (angiotensin precursor; F_(1,33)_=8.272, *p*=0.007) mRNA in male mice (*p*=0.034). CUS increased astrocyte expression of *Aqp4* (F_(1,34)_=11.36, *p*=0.002) and the vascular stabilization gene *Angpt1* (F_(1,34)_=16.58, *p*<0.001) in male mice, yet decreased these transcripts in females. A number of basal sex differences were also detected in astrocytes in the frontal cortex, including reduced levels of *S100b*- and heightened levels of *Mertk*, *Vegf* (angiogenic factor; F_(1,34)_=9.087, *p*=0.005), *Aqp4*, and *Angpt1* transcript in female as compared to male mice (planned comparison, *p*<0.05). Collectively, these results indicate divergent, sex-specific stress effects on astrocyte gene expression in the frontal cortex, with increased expression of genes involved with vascular interactions in males (Fig 2B).

**Figure 2.**
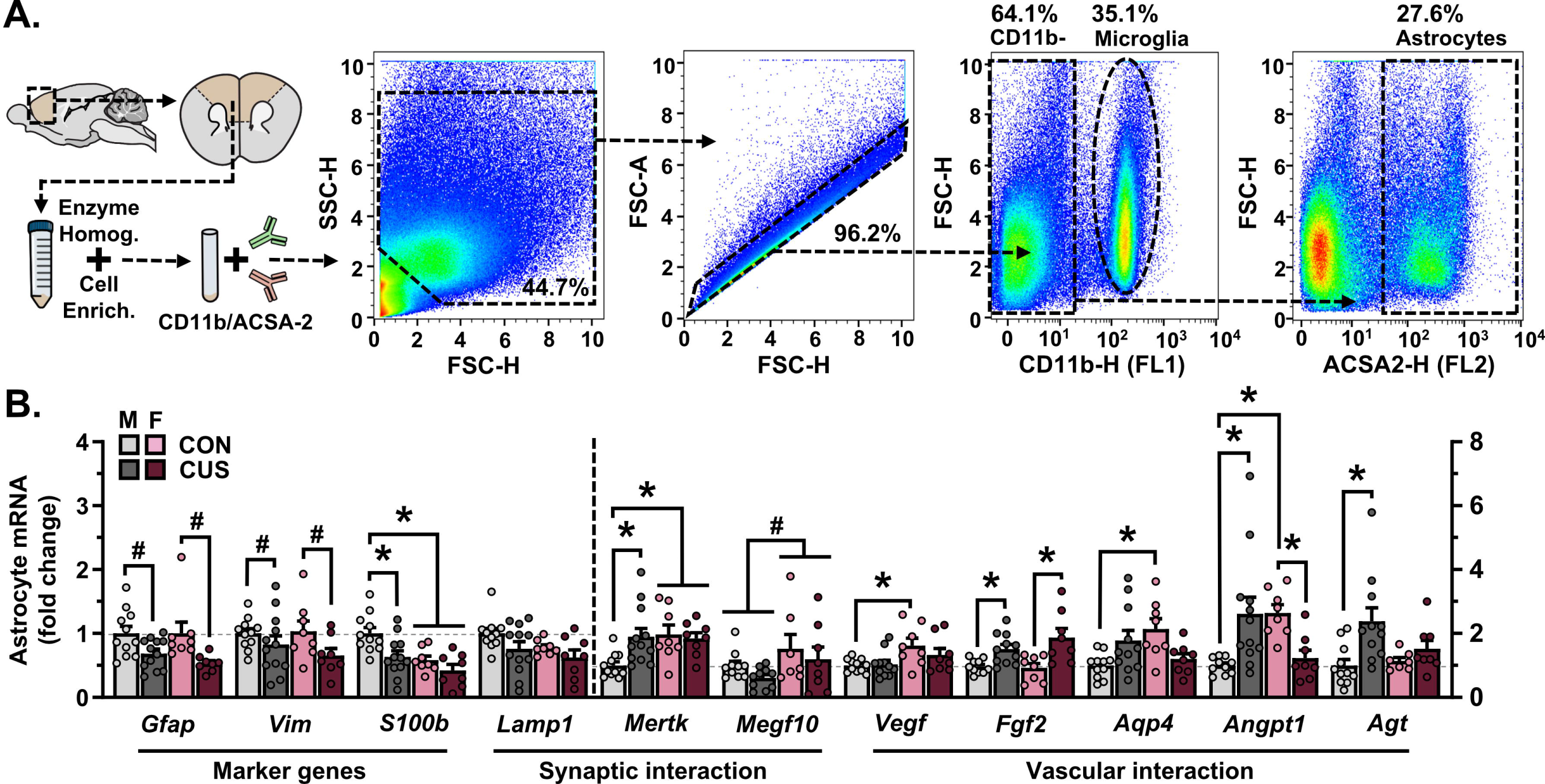
Chronic unpredictable stress (14-days) differentially alters the molecular phenotype of astrocytes in the prefrontal cortex in males and females. **A)** A prefrontal cortex-enriched portion of the brain was dissected out and homogenized. Cells were then stained for CD11b and ACSA2 using fluorescently conjugated antibodies and characterized using flow cytometry. CD11b-/ACSA2+ astrocytes were collected using FACS (proportion of gated events relative to the total number of events in each plot is indicated). Representative gating strategy is shown. **B)** Normalized astrocyte expression of marker genes (*Gfap*, *Vim*, *S100b*), synaptic interaction factors (*Lamp1*, *Mertk*, *Megf10*), and vascular interaction factors (*Vegf*, *Bfgf*, *Aqp4*, *Angpt1*, *Agt*) in the frontal cortex. Expression of *Gfap*, *Vim*, *S100b*, and *Lamp1* is reflected on the left y-axis, all other genes are reflected on the right y-axis. Bars represent mean ± S.E.M. ^#^ p<0.05 main effect (no specific contrasts detected). * p<0.05 planned comparison indicated (Sidak’s test).

### Stress-induced atrophy of astrocytes in the PFC is associated with loss of neurovascular AQP4 and working memory dysfunction in male -but not female-mice

Blood-brain barrier permeability and astrocyte-neurovascular interactions were examined in another subset of mice (d15). For these experiments, Alexa Fluor 488-conjugated dextran was injected intravenously (30 min circulation) followed by tomato lectin (5 min circulation). Mice were then perfused, and brains were extracted, sectioned, stained, and visualized using confocal microscopy (Fig 3A). Neither biological sex nor CUS affected the density of ALDH1L1+ astrocytes in the PFC (Fig 3B, Fig S2). Nonetheless, the amount of total GFAP+ astrocytic material (F_(1,27)_=8.629, *p*=0.007; Fig 3C) and blood vessel-associated GFAP (F_(1,23)_=5.223, *p*=0.032; Fig 3D) differed between males and females. CUS diminished the amount of total (*p*=0.002)- and blood vessel-associated GFAP in male mice only (*p*=0.005). Levels of AQP4+ material-(F_(1,26)_=4.290, *p*=0.048; Fig S2) and astrocyte AQP4-blood vessel coverage (F_(1,24)_=27.51, *p*<0.0001; Fig 3E) were reduced in females as compared to males. Exposure to CUS decreased AQP4-rich astrocyte endfoot coverage of blood vessels in the PFC in males (*p*=0.009), yet increased coverage in females (*p*=0.002). CUS increased blood vessel (Tomato lectin+) area in male mice (F_(1,25)_=14.56, *p*<0.001; planned comparison: *p*=0.013; Fig 3F), but decreased this in females (*p*=0.049). In contrast to the PFC, neither sex-nor stress affected astrocyte GFAP+ area, AQP4 coverage of blood vessels, or vessel area in the S1J subregion of the somatosensory cortex (SSCTX, Fig S3). Analysis of BBB integrity in the PFC revealed an increased number of dextran extravasation sites (F_(1,25)_=10.32, *p*=0.004) and heightened levels of dextran fluorescence (F_(1,25)_=4.283, *p*=0.049) in female as compared to male mice. CUS increased the number (F_(1,25)_=7.926, *p*=0.009)- and fluorescence intensity (F_(1,25)_=6.035, *p*=0.021) of dextran extravasation sites in both sexes. Remodeling of GFAP+ astrocytic structures was associated with loss of vascular AQP4 coverage in the PFC (r=0.580, p=0.030) and impairment in the TOR (r=0.509, p=0.044) in male -but not female-mice. Interestingly, astrocytic structures (GFAP and AQP4 coverage) were not associated with blood vessel area or extravasation in either sex (Fig S2). Our results indicate that CUS leads to astrocyte atrophy (GFAP+ structural loss) and diminished astrocytic coverage of blood vessels within the PFC of male mice. Likewise, remodeling of astrocytic structures is associated with impairment in PFC-mediated behavior in males only.

**Figure 3.**
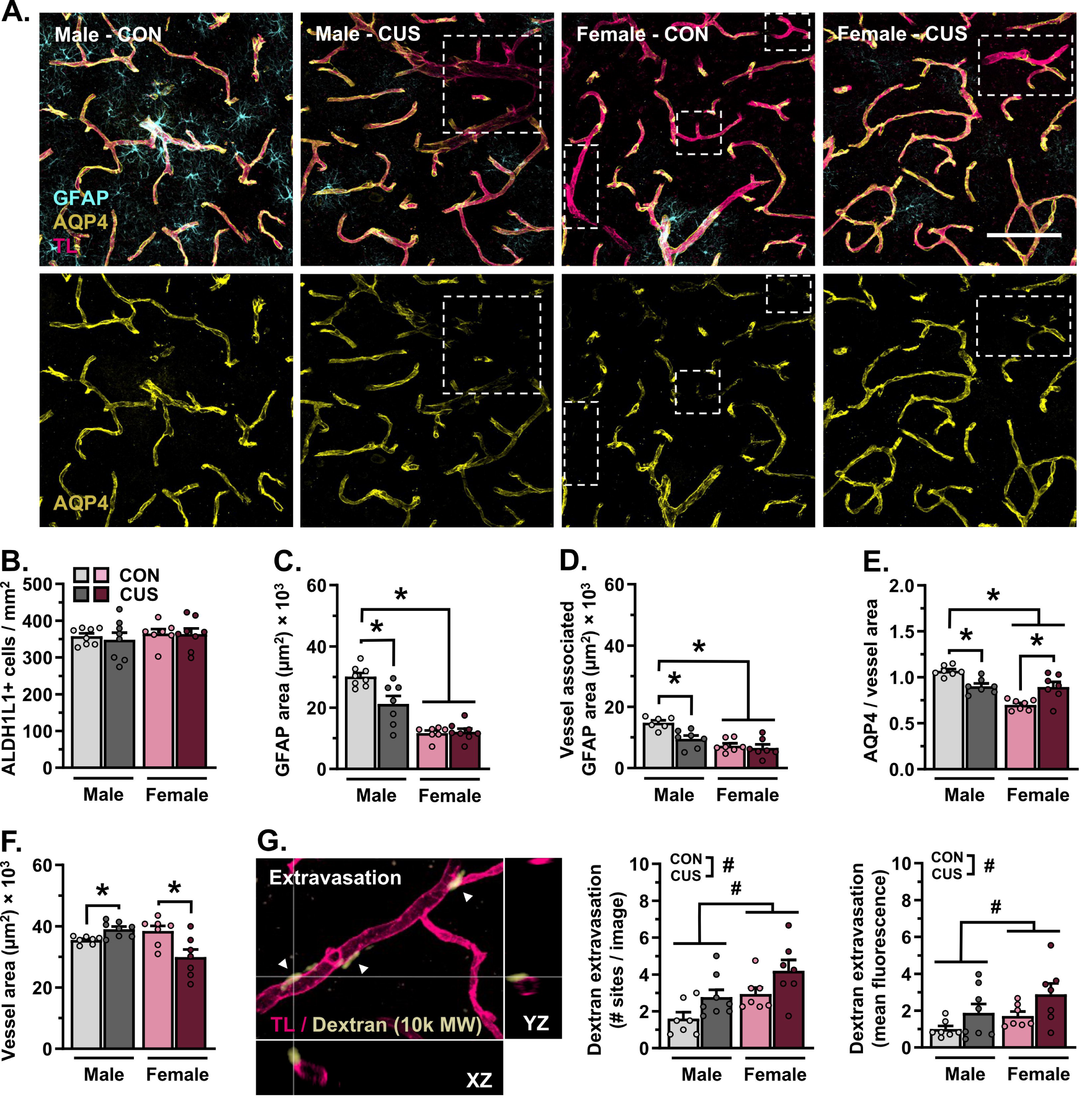
Chronic stress (14-days) leads to astrocyte atrophy and vascular-associated aquaporin-4 loss in the prefrontal cortex in male, but not female, mice. **A)** Representative images of vascular tomato lectin (TL) staining (magenta) and immunohistology for astrocyte GFAP (cyan) and AQP4 (yellow) in the PFC (20× + zoom, scale bar = 100 µm). Dashed quadrilaterals indicate TL+ vessel segments with little-to no AQP4 coverage. **B)** Number of ALDH1L1+ astrocytes in the PFC per mm^2^. **C)** Area of GFAP+ material in the PFC. **D)** Area of GFAP+ material associated with blood vessels in the PFC. **E)** Area of astrocyte AQP4+ material relative to vessel area. **F)** Area of tomato lectin+ vessels in the PFC. **G)** Left: Representative image of Alexa Fluor 488-conjugated dextran (10,000 MW, pale yellow) and vascular tomato lectin (magenta) in the PFC. Image features three extravasation sites, with one site highlighted in XZ-YZ dimensions. Middle: Number of dextran extravasation sites in the PFC per image. Right: Mean fluorescence intensity of dextran extravasation sites in the PFC (normalized to control males). Bars represent mean ± S.E.M. ^#^ p<0.05 main effect (no specific contrasts detected). * p<0.05 planned comparison indicated (Sidak’s test).

### Knockdown of AQP4 in the PFC leads to behavioral stress susceptibility in males and disrupts the molecular phenotype of PFC astrocytes

In subsequent experiments, AQP4 was knocked down in the PFC of male mice using an AAV5-*Aqp4*-shRNA construct; control animals received infusions of AAV5-scrambled-shRNA (Fig 4A). Infusion of AAV5-*Aqp4*-shRNA into the PFC robustly infected astrocytes (Fig 4B) and reduced *Aqp4* expression in the PFC but not the adjacent S1J subregion of the SSCTX (F_(1,12)_=5.164, *p*=0.042; Fig 4C). Following surgical recovery (3 weeks), mice were either left unstressed or exposed to 7 days of CUS (sub-CUS). This shortened stress paradigm has limited effects on behavior. All mice were assessed in the OFT (d5), TOR (d6), and a T-maze task (d7). Neither *Aqp4* knockdown nor sub-CUS affected locomotion in the OFT (Fig 4D). There was an interaction between *Aqp4* knockdown and sub-CUS on the amount of time mice spent exploring the OFT center zone (F_(1,49)_=4.874, *p*=0.032; Fig 4E), however no specific contrasts were detected. In the TOR, mice spent a similar amount of time engaging with objects, regardless of *Aqp4* knockdown or sub-CUS exposure (Fig 4F). Despite this, only mice with *Aqp4* knockdown in the PFC and sub-CUS showed deficits in object discrimination (F_(1,55)_=7.634, *p*=0.008; Fig 4G). In the T-maze, mice successfully discriminated between a previously explored- and novel arm (scrambled-shRNA: F_(1,56)_=38.83, *p*<0.0001, *Aqp4*-shRNA: F_(1,56)_=27.55, *p*<0.0001; Fig 4H). However, depletion of *Aqp4* reduced the number of alternations made during the test phase (F_(1,53)_=5.016, *p*=0.029; Fig 4I). Exposure to sub-CUS enhanced spontaneous alternation behavior in mice that received the AAV5-scrambled-shRNA construct (F_(1,54)_=4.155, *p*=0.046, planned comparison: *p*=0.038); this was not observed in mice with *Aqp4* knockdown (Fig 4J). Collectively, these results suggest that loss of AQP4 in PFC astrocytes leads to stress susceptibility in cognitive tasks.

**Figure 4.**
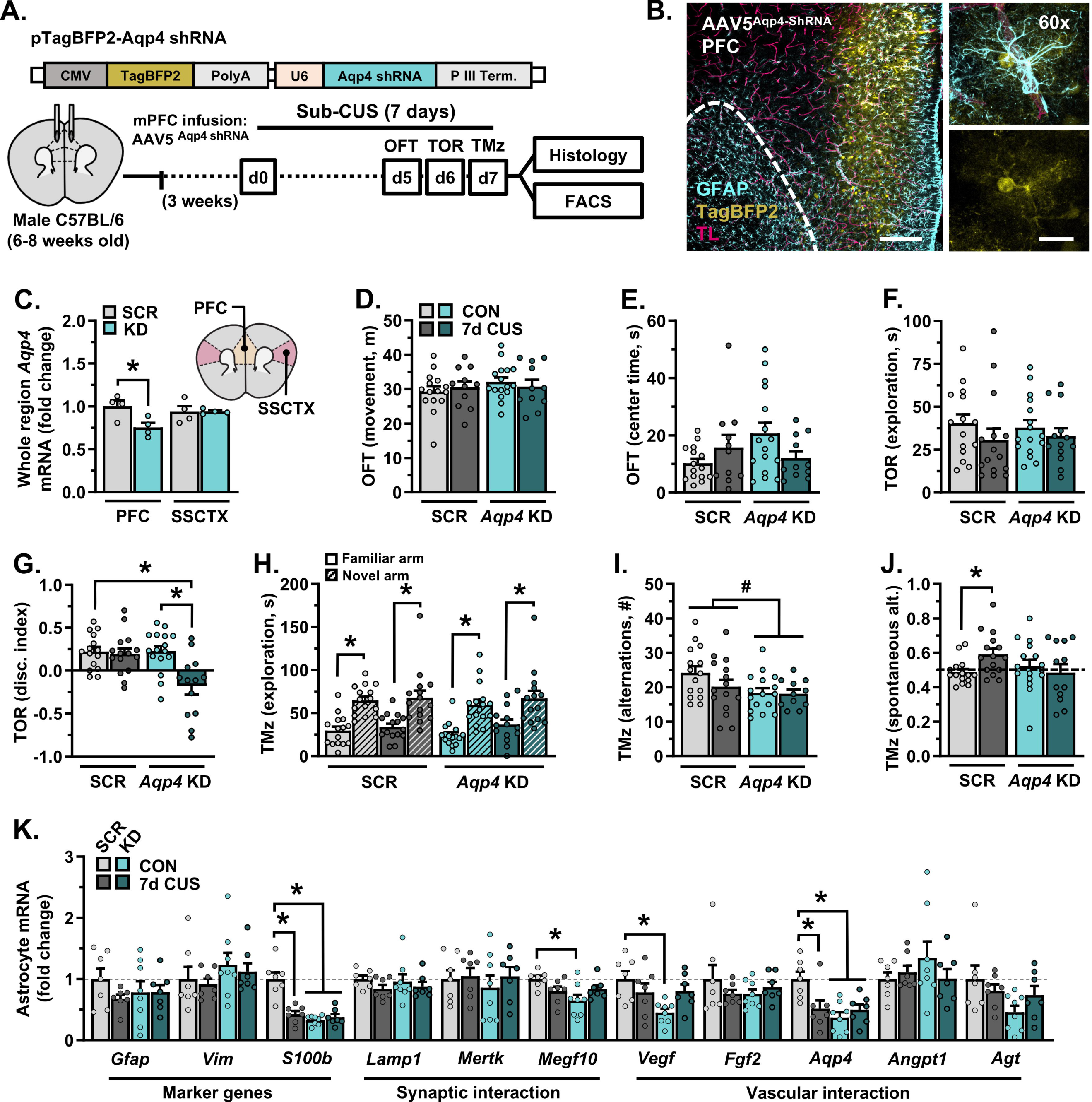
Knockdown of aquaporin-4 in the prefrontal cortex shifts astrocyte phenotype and induces behavioral susceptibility to a sub-chronic stressor (7-days) **A)** Adult male mice received infusions of either AAV5-scrambled-shRNA or AAV5-*Aqp4*-shRNA into the PFC. Following three weeks of recovery, animals were exposed to 7-days of sub-CUS or were left unstressed. All animals were subjected to behavioral testing (n = 11-16/group). In one cohort, mice received intravenous injections of tomato lectin, after which brains were processed for confocal microscopy (n = 6-8/group). In a separate cohort, astrocytes were isolated via FACS and astrocyte gene expression was analyzed (n = 7-8/group). **B)** Representative images of AAV5-*Aqp4*-ShRNA reporter (BFP, yellow) in the PFC. Astrocyte GFAP is shown in cyan, tomato lectin+ vessels are shown in magenta. Dotted line delineates barrier between the forceps minor and PFC. Left image: 10×, scale bar = 200 µm. Right image panel: 60×, scale bar = 20 µm. **C)** Normalized expression of *Aqp4* in dissected PFC or SSCTX (S1J subregion). **D)** Total amount of distance traveled in the open field test (OFT). **E)** Time spent in the center during the OFT. **F)** Total amount of time spent exploring objects in the test phase of the temporal object recognition task (TOR). **G)** Discrimination index in the TOR. **H)** Time spent exploring both the novel and familiar arm in the T-maze task. **I)** Total number of alternations in the T-maze task. **J)** Proportion of spontaneous alternations in the T-maze task. **K)** Normalized astrocyte expression of marker genes (*Gfap*, *Vim*, *S100b*), synaptic interaction factors (*Lamp1*, *Mertk*, *Megf10*), and vascular interaction factors (*Vegf*, *Bfgf*, *Aqp4*, *Angpt1*, *Agt*) in the frontal cortex. Bars represent mean ± S.E.M. ^#^ p<0.05 main effect (no specific contrasts detected). * p<0.05 planned comparison indicated (Sidak’s test).

Gene expression in PFC astrocytes was then characterized in a subset of mice using the same protocol as described above (d8). Despite having no effect on behavior, exposure to sub-CUS altered the molecular phenotype of astrocytes in the frontal cortex in animals with AAV5-scrambled-shRNA (Fig 4K). This included decreased levels of *S100b* (F_(1,25)_=22.56, *p*<0.0001) and *Aqp4* transcripts (F_(1,25)_=7.960, *p*=0.009). Virally mediated knockdown of *Aqp4* also reduced expression levels of *S100b* and *Aqp4*, alongside *Megf10* (F_(1,25)_=6.647, *p*=0.016) and *Vegf* (F_(1,25)_=6.311, *p*=0.019) in the frontal cortex. Together, these data indicate that transcripts reflecting the phenotype of astrocytes shift prior to the emergence of stress-induced deficits in behavior, and that depletion of *Aqp4* in PFC astrocytes is sufficient to disrupt markers of astrocyte identity (i.e., *S100b*) and function (i.e., *Megf10, Vegf*).

### AQP4 knockdown in the PFC and sub-CUS induce astrocyte dystrophy

A separate cohort of mice received intravascular injections of tomato lectin prior to perfusion and immunostaining (d8), with subsequent analyses focused on astrocyte-blood vessel interaction in the PFC (Fig 5A). *Aqp4* knockdown reduced the density ALDH1L1+ astrocytes in the PFC (F_(1,26)_=9.022, *p*=0.006; Fig 5B). Sub-CUS had no effect on the number of PFC astrocytes in mice that received infusions of AAV5-scrambled-shRNA, yet reduced the total amount of GFAP+ material (F_(1,26)_=29.24, *p*<0.0001; Fig 5C), blood vessel-associated GFAP+ material (F_(1,24)_=31.25, *p*<0.0001; Fig 5D), AQP4+ material (F_(1,26)_=4.599, *p*=0.042; Fig 5E), and astrocyte AQP4-blood vessel coverage (F_(1,24)_=8.250, *p*=0.008; Fig 5F) in this region. Similar to sub-CUS, shRNA-mediated loss of *Aqp4* diminished the amount of GFAP+ material, blood vessel-associated GFAP, AQP4+ material, and astrocyte AQP4-rich endfoot coverage of blood vessels in the PFC. Decreased expression of *Aqp4* slightly increased vessel area in the PFC (F_(1,25)_=4.420, *p*=0.046; Fig 5G).

**Figure 5.**
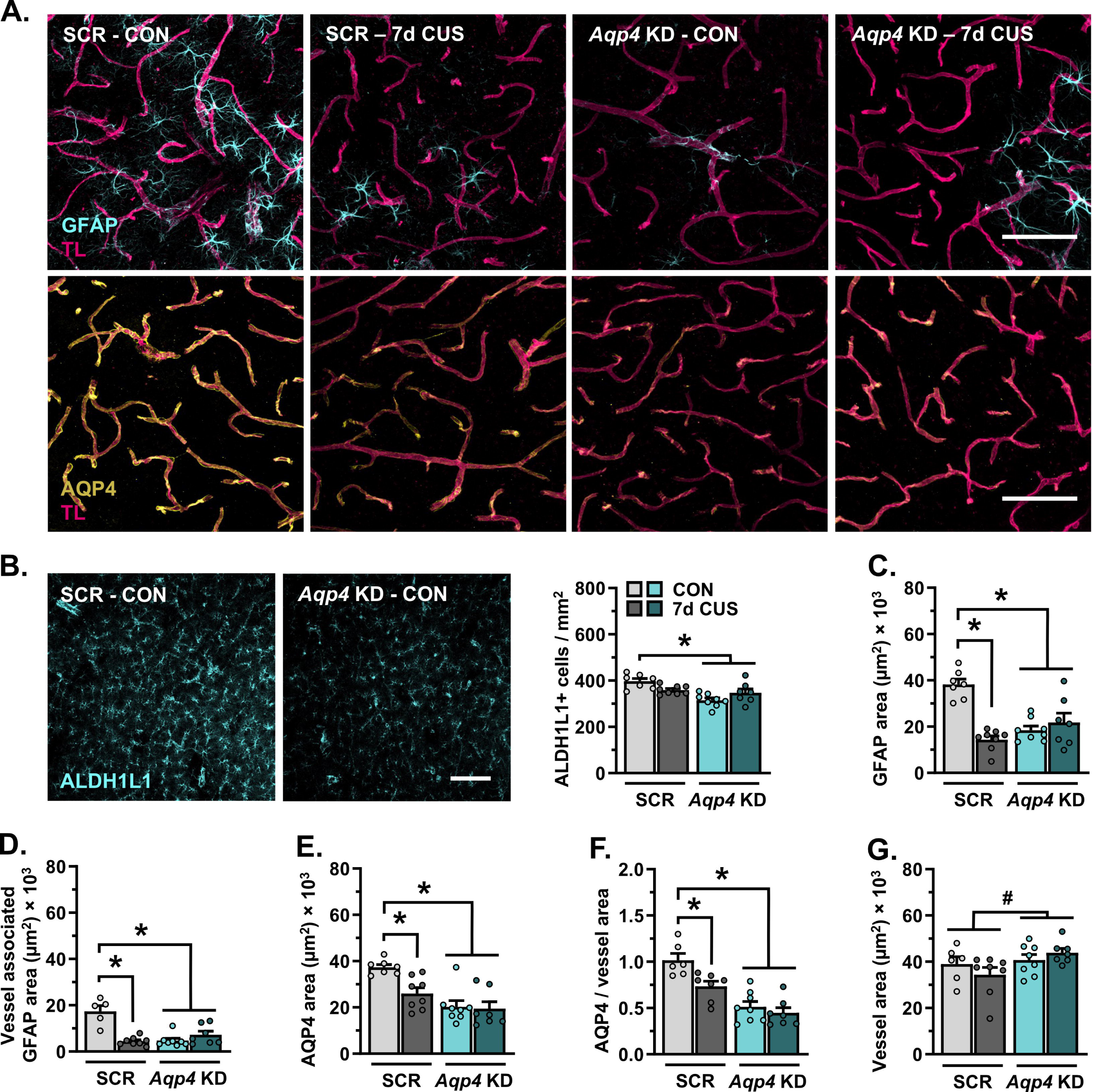
Astrocytes dynamically respond to sub-chronic stress, knockdown of aquaporin-4 leads to astrocyte dystrophy in the prefrontal cortex. **A)** Representative images of vascular tomato lectin (magenta) and immunohistology for astrocyte GFAP (cyan) and AQP4 (yellow) in the PFC (20×, scale bar = 100 µm). **B)** Left: Representative images of immunohistology for ALDH1L1 in the PFC (20×, scale bar = 100 µm). Right: Number of ALDH1L1+ astrocytes in the PFC per mm^2^. **C)** Area of GFAP+ material in the PFC. **D)** Area of GFAP+ material associated with blood vessels in the PFC. **E)** Total area of AQP4+ material in the PFC. **F)** Area of astrocyte AQP4+ material relative to vessel area. **G)** Area of tomato lectin+ vessels in the PFC. ^#^ p<0.05 main effect (no specific contrasts detected). * p<0.05 planned comparison indicated (Sidak’s test).

## Discussion

Postmortem studies indicate that disruptions in astrocyte phenotype and function, including loss of perivascular processes and blood vessel associated AQP4-rich endfeet, is observed in individuals diagnosed with MDD [13]. Likewise, astrocyte dystrophy in the PFC is evident in rodent models of chronic stress [15], with separate studies noting impaired BBB function in various stress-susceptible brain regions [28, 42]. Here, we extend these findings to show that chronic stress remodels astrocytes at the PFC neurovascular interface in male – but not female – mice. Our data further demonstrate a role for AQP4 in regulating not only astrocyte gene expression and structure in the PFC, but susceptibility to stress-induced deficits in working memory function in males. Alongside these findings, we uncovered several basal sex differences in astrocyte phenotype, blood vessel morphology, and neurovascular integrity in the PFC that may underlie sex-specific responses to stress.

### Chronic stress disrupts astrocyte-blood vessel interaction in the PFC in males, loss of prefrontal AQP4 induces behavioral stress susceptibility

As previously shown, exposure to CUS led to astrocyte dystrophy in the PFC in male mice [22, 43]. This included retraction of perivascular astrocytic processes and AQP4-rich endfeet. Our findings also indicate increased blood vessel area and small molecule extravasation across the prefrontal BBB. Initially, we thought that stress-induced reductions in AQP4 may underlie these alterations in neurovascular structure and permeability. However, our data do not support this hypothesis (neither levels of GFAP-nor AQP4 correlated with vessel area or dextran extravasation). Indeed, prior studies indicate that chronic stress causes degradation of endothelial tight junctions and loss of intrinsic factors critical in blood vessel coupling, including claudin-5 [26, 28, 44]. Genetic knockdown of claudin-5 alone impairs the prefrontal BBB leading to anhedonia-like behaviors [28]. How astrocytes respond to loss of claudin-5 remains unknown. Considering this, it is possible that stress-induced permeabilization of the BBB occurs in tandem with perivascular astrocyte reorganization. It is likely that these neurobiological consequences together contribute to stress-induced alterations in PFC function and behavior.

Alongside retraction from blood vessels, chronic stress caused prefrontal astrocytes to engage various molecular pathways indicative of neurovascular distress. This included upregulation of astrocytic *Mertk*, *Fgf2*, *Angpt1*, and *Agt* expression. These factors are all linked with blood vessel maintenance, growth, and function, and perhaps reflect a need for astrocyte-mediated repair [45]. Additional studies indicate that similar angiogenesis-associated pathways are upregulated in whole tissue samples- and brain endothelial cells during chronic stress [24, 46]. Again, these lines of evidence support the notion that astrocytes are responding to-but not necessarily driving neurovascular dysfunction.

Beyond BBB permeability, AQP4 depletion in PFC astrocytes induced behavioral susceptibility to stress, as reflected by a deficit in temporal object recognition following subthreshold-CUS. Knockdown of AQP4 also led to astrocyte atrophy and diminished expression of astrocytic factors implicated in blood vessel interaction in the PFC. If not through BBB modulation, there are other pathways by which loss of astrocyte AQP4 may impact brain function and behavior. For instance, recent studies indicate that chronic stress impairs glymphatic influx across the neocortex [9, 18, 19]. Glymphatic influx allows for the removal of waste products from the brain through AQP4-dependent blood vessel efflux [11, 47, 48]. This is tightly coupled to neuronal activity [47, 48]. As such, it is conceivable that loss of AQP4 impaired glymphatic flux in the PFC. While viral knockdown of AQP4 had little effect on cognition at baseline, in combination with stress – a paradigm which provokes heightened prefrontal activity – it is possible that loss of AQP4 exacerbated waste buildup in the PFC, impairing both neuronal function and working memory [49, 50]. Follow-up studies will be needed to delineate the consequences of stress induced AQP4 loss on prefrontal glymphatic dynamics and neuronal health.

### Basal sex differences in- and divergent stress effects on-astrocyte phenotype and blood vessel coverage in the PFC

In contrast to males, stress decreased blood vessel area in the PFC and increased AQP4 coverage of the neurovasculature in females. This was accompanied by a unique shift in the molecular phenotype of prefrontal astrocytes, reflecting both up- and downregulation of pathways important in blood vessel maintenance. These divergent stress effects may arise from pronounced sex differences in astrocyte state and neurovascular engagement in the PFC. For instance, basal levels of astrocytic GFAP+ material are diminished in the PFC in females as compared to males [22, 43]. This was not observed in the SSCTX, suggesting region specific sex differences in astrocyte phenotype [51]. Likewise, we found increased levels of *Mertk*, *Vegf*, *Aqp4*, and *Angpt1* mRNA in prefrontal astrocytes from females as compared to males. Interestingly, this broad upregulation of neurovascular-associated transcripts in astrocytes occurred alongside reduced coverage of blood vessels by astrocytic processes and heightened dextran extravasation in female mice [52]. Akin to our findings in stressed males, this would suggest that astrocytes are receptive to blood vessel-associated damage signals in the PFC and are responding to these signals through the upregulation of factors important in angiogenesis and repair.

These pronounced sex differences are likely mediated by a combination of developmental influences and gonadal hormone-dependent mechanisms. For instance, recent studies indicate distinct transcriptional and morphological patterns of astrocyte development in the cortex in male and female mice [51]. These developmental patterns are dependent on early life exposure to estradiol [51, 53]. Similarly, we’ve shown that stress-induced atrophy of astrocytes in the PFC is mediated, in part, by gonadal hormones in adult males, and that estradiol can modulate stress-induced morphological changes in astrocytes in both sexes [43]. Astrocytes express androgen and estrogen receptors, therefore it’s plausible that gonadal hormones directly act on-, regulate-, and shape the function of these cells [33, 54, 55]. It is also possible that other cell types interacting with astrocytes, including neurons, additional glia, and endothelial cells, indirectly mediate the effects of gonadal hormones on astrocytic structures. Future studies examining the role of astrocyte androgen and estrogen receptors in stress, and their respective influences on sex differences in astrocyte-neurovascular engagement are warranted.

### Conclusion

The present study uncovered several striking sex differences in astrocyte transcription, astrocyte-blood vessel interaction, and neurovascular integrity in the PFC. Of note, chronic stress caused astrocyte atrophy and diminished AQP4-endfoot coverage of blood vessels that was associated with behavioral and cognitive consequences in male mice only. Mechanistic studies indicate that viral-mediated AQP4 knockdown in PFC astrocytes is sufficient to alter astrocyte phenotype and structure, leading to behavioral susceptibility in a subthreshold stress model. These preclinical findings are similar to those of recent transcriptomic reports examining the PFC in men and women with MDD. Though few, these studies indicate reduced levels of astrocyte-associated genes and astrocyte *Aqp4* transcript in the PFC of men but not women with MDD [56–58]. Another study also found increased levels of *Aqp4*-antisense RNA 1, which downregulates AQP4 mRNA translation, in the PFC of men with MDD [59]. Collectively, these findings support a sex-specific role for prefrontal astrocytes and AQP4-coverage of blood vessels in chronic psychological stress and MDD.

## Acknowledgments

This work was supported by funding from the National Institutes of Health (F32 MH123051 to JLB; R01 MH123545 to ESW) and generous support from the University of Cincinnati College of Medicine. We would like to thank the users of Scidraw for providing free use of the following graphics (https://doi.org/10.5281/zenodo.3925917)

---

**Supplementary Figure 1. Chronic stress reduces animal weight gain. A)** Weight change in mice subjected to CUS or left unstressed (14-days). Stress differentially reduced weight gain in male and female mice (F_(1,75)_=8.148 *p*=0.005). Planned comparisons indicate a significant reduction in weight gain in stressed males (*p*<0.0001) and a trend toward this in stressed females (*p*=0.089). **B)** Weight change in mice subjected to sub-CUS or left unstressed (7-days). Exposure to sub-CUS reduced weight gain in mice, regardless of *Aqp4* knockdown in the PFC (F_(1,26)_=41.98 *p*<0.001). Bars represent mean ± S.E.M. * p<0.05 comparison indicated (Sidak’s test).

**Supplementary Figure 2. Loss of astrocytic structures is associated with diminished aquaporin-4 blood vessel coverage in the prefrontal cortex and working memory impairment in male mice. A)** Representative images of ALDH1L (Cyan) and GFAP (yellow) immunohistology in the PFC (20×, scale bar = 50 µm. **B)** Total area of AQP4+ material in the PFC. **C)** Linear association between GFAP+ area and AQP4 blood vessel coverage in the PFC. **D)** Linear association between GFAP+ area in the PFC and discrimination in the TOR. **E)** Linear association between GFAP+ area and the number of dextran extravasation sites in the PFC. **F)** Linear association between AQP4 blood vessel coverage and the number of dextran extravasation sites in the PFC. Bars represent mean ± S.E.M. * p<0.05 comparison indicated (Sidak’s test or Pearson’s correlation).

**Supplementary Figure 3. Analysis of astrocyte coverage and blood vessel area in the somatosensory cortex. A)** Schematic of the S1J subregion of the SSCTX. Analysis of this region occurred in the same animals as presented in Figure 3. **B)** Area of GFAP+ material in the SSCTX. **C)** Total area of AQP4+ material in the SSCTX. **D)** Area of astrocyte AQP4+ material relative to vessel area. **E)** Area of tomato lectin+ vessels in the SSCTX. Bars represent mean ± S.E.M.

