## Supplementary figures and images for "Stress-induced dysfunction of neurovascular astrocytes contributes to sex-specific behavioral deficits"

### Supplemental Figure 1

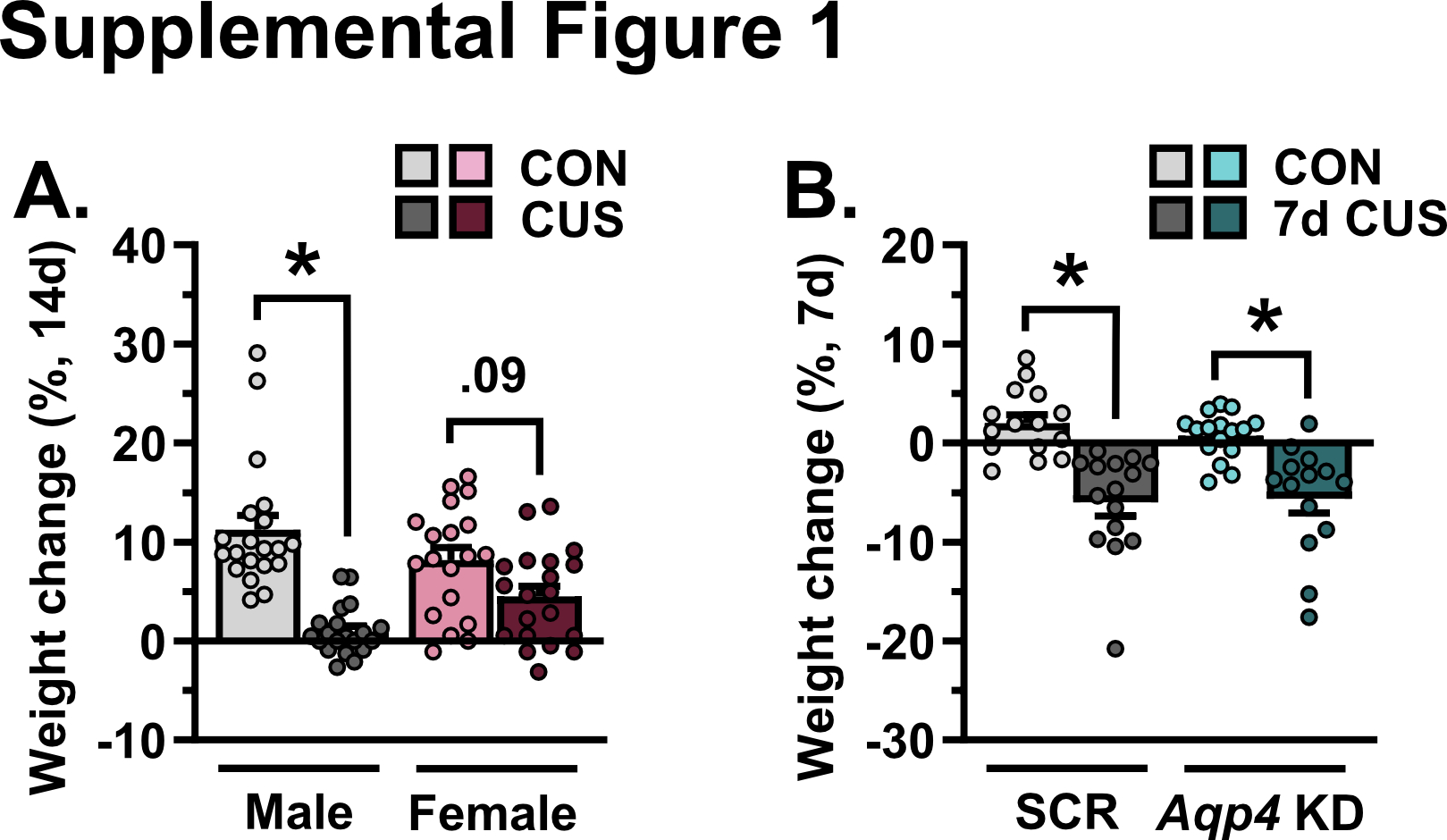

### Supplemental Figure 2

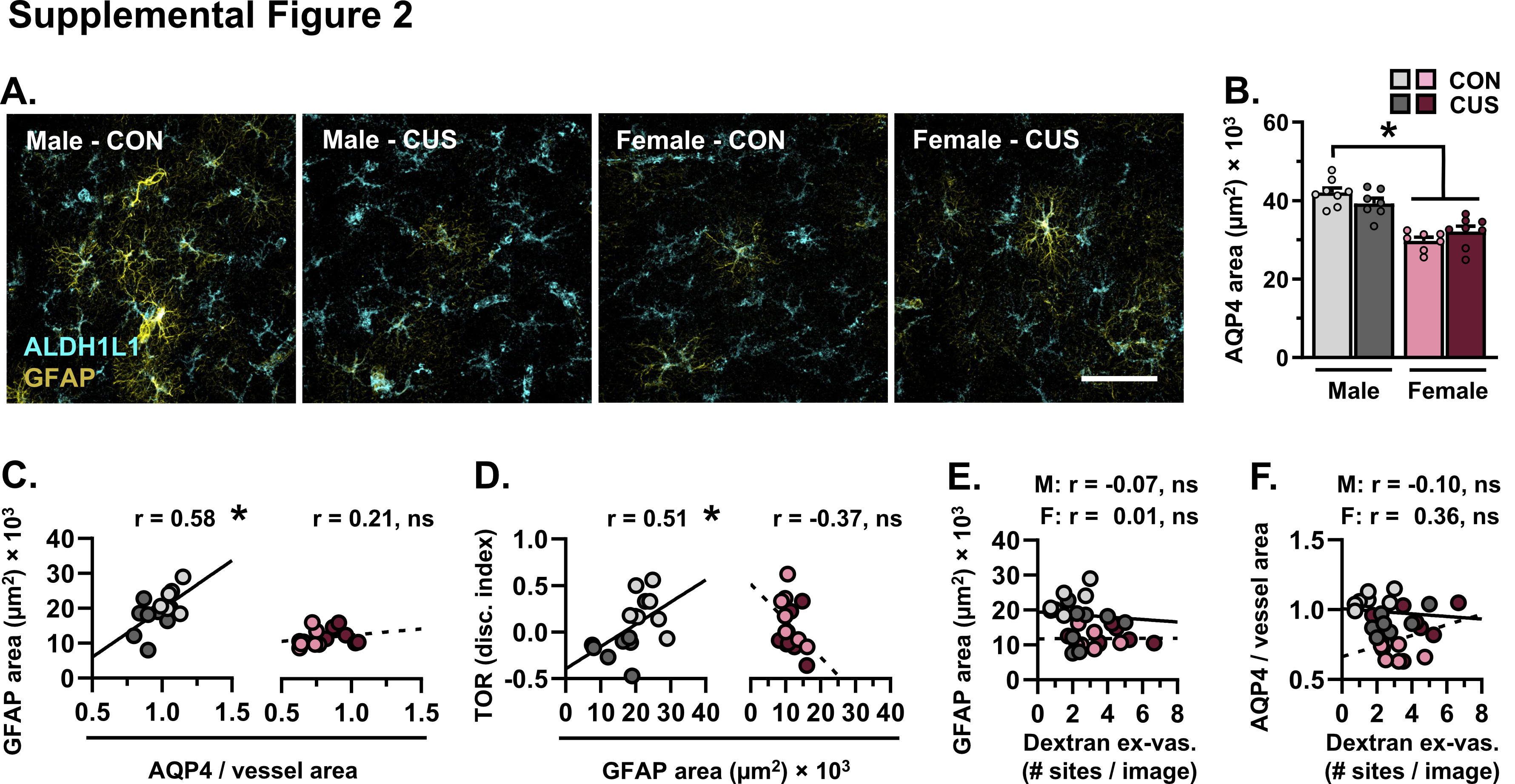

### Supplemental Figure 3

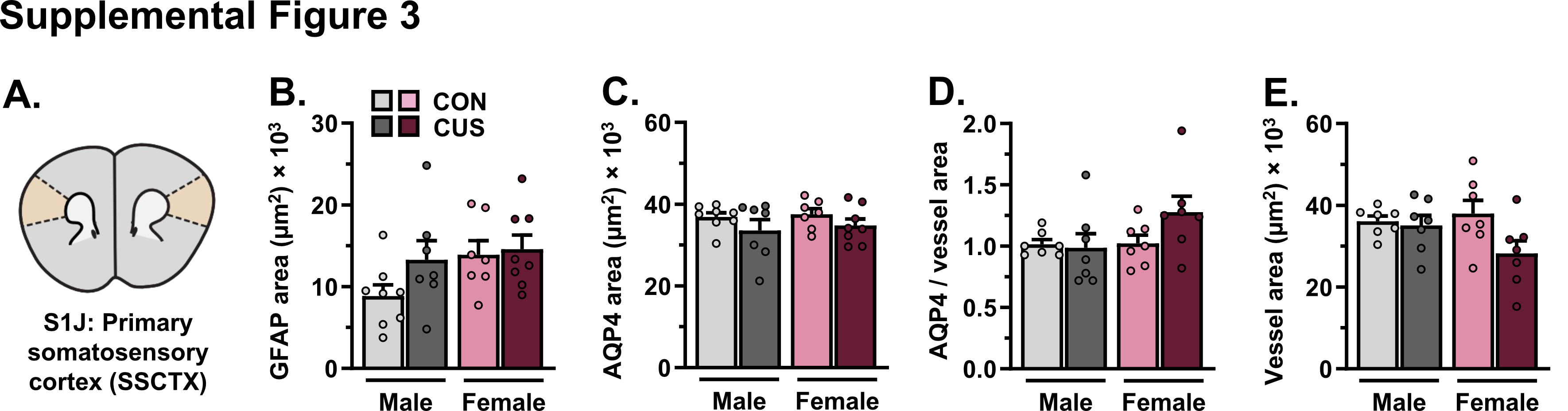
